# An epigenetic switch in vascular phenotype augments anti-tumor immunity

**DOI:** 10.64898/2025.12.05.692381

**Authors:** Dae Joong Kim, Mitchell McGinty, Swetha Anandh, Caroline Riedstra, Yuvraj Sethi, Melanie R. Rutkowski, Andrew C. Dudley

**Affiliations:** Department of Microbiology, Immunology, and Cancer Biology, The University of Virginia, Charlottesville, VA 22908, USA; The UVA Comprehensive Cancer Center, The University of Virginia, Charlottesville, VA 22908, USA

**Keywords:** Tumor endothelial cells, DNA methyltransferase, tumor immune microenvironment, angiogenesis, immunotherapy, melanoma

## Abstract

The abnormal tumor vasculature can present a barrier to the infiltration of anti-tumor immune cells, which impairs immune surveillance and response to immunotherapy. Here, we show that targeting the epigenetic factor DNA methyltransferase 1 in endothelial cells (ECs) reduces angiogenesis while imparting profound changes to the tumor immune microenvironment (TIME), including increased proportions of CD4^+^ memory T-cells and NK cells. Depleting CD4^+^ T-cells, or blocking lymphocyte egress from the lymph nodes with FTY720, rescues tumor growth in mice with conditional deletion of Dnmt1 in ECs (*Dnmt1*^iECKO^) and dramatically shortens overall survival, whereas NK cells are dispensable. Tumors implanted in *Dnmt1*^iECKO^ mice show reduced vascular branching, elevated expression of Vcam1, increased vessel-associated T-cells, and a shift in vascular specification including increased proportions of immune-permissive post-capillary venules (PCVs) and interferon-stimulated ECs (IFN-ECs). Deleting Dnmt1 in EC cultures strikingly potentiates responses to combinations of IFNγ and TNFα and, notably, up-regulates important T-cell co-stimulatory molecules for memory CD4^+^ T-cells, including *Icosl*, *Cd40*, and *Tnfsf4*. Finally, immune checkpoint blockade (ICB) administered to *Dnmt1*^iECKO^ mice with experimental melanoma lung metastasis reduces tumor burden, with some mice showing tumor eradication. Our findings identify endothelial Dnmt1 as a key regulator of vascular-mediated anti-tumor immunity, providing a rationale for integrating epigenetic modulation of the vasculature with cancer immunotherapy regimens.

## INTRODUCTION

Tumor-associated blood vessels are a point of entry for anti-tumor immune cells into the tumor immune microenvironment (TIME) (1). However, abnormalities in the vasculature, including disruptions in blood flow, aberrant or reduced expression of cell adhesion molecules (CAMs) that are critical for T-cell entry, or elevated expression of PD-L1 consort to dampen a robust anti-tumor immune response (2–4). In these tumors, the absence of effector T-cells or T-cell exhaustion, coupled with insufficient antigen presentation, allows cancer cells to escape elimination (5). Recently, immune checkpoint blockade (ICB) has emerged as a promising treatment option to reinvigorate anti-tumor immunity, resulting in improved survival for many patients; this is especially true in advanced-stage melanomas with high tumor mutational burden (TMB) and in certain hypermutant colorectal cancers with defective mismatch repair (6, 7). However, not all of these patients respond to ICB, and resistance mechanisms can develop, indicating there is potential to improve the efficacy of ICB by either customizing dose regimens or by adding a secondary drug. For example, drugs that target the epigenome are frequently combined with ICB and have proven safe and effective in pre-clinical models and clinical trials (8, 9).

Similarly, combinations of ICB with anti-angiogenic (AA) therapies are beneficial in mice and humans, which has motivated several new clinical trials across a spectrum of cancer types (10–13). Multiple mechanisms could account for the beneficial effect of combining AA therapy with ICB. For example, it is suggested that judicious doses of AA therapies normalize the tumor vasculature and derepress the expression of CAMs and chemokines that are essential for T-cell entry (14, 15). It is also possible that AA therapies reprogram tumor blood vessels by down-regulating the expression of endothelial cell (EC)-derived chemokines or other factors that recruit immunosuppressive myeloid cells or regulatory T cells (16); thus, AA therapies have the potential to promote large-scale changes in the TIME by changing the function or phenotype of the tumor vasculature. A good example is the therapeutic induction of high endothelial venules (HEVs), which are specialized for lymphocyte trafficking (17). As a corollary, combining anti-VEGF therapy with lymphotoxin beta receptor (LTβR) agonists, or ICB, promotes the de novo formation of HEVs or HEV-like vessels in preclinical models, suggesting that vascular specification in tumors can be strategically tailored to promote T-lymphocyte entry (18–21). These types of strategies could be especially beneficial for CAR-T therapies, which often fail due to the poor homing to or extravasation of engineered T-cells from the vasculature (22).

Other avenues for targeting tumor blood vessels to augment anti-tumor immunity, or boost ICB efficacy, include the activation of inflammatory pathways (e.g. cGAS/Sting agonism), irradiation/chemotherapy to reverse tumor EC anergy, metabolic rewiring of tumor EC functions, and epigenetic or transcription-factor driven reprogramming of tumor ECs [reviewed in (3)]. A commonality in many of these approaches is the upregulation of CAMs (e.g. VCAM1 or ICAM1) and Th1 chemokines (e.g. Cxcl9/10) that are displayed on the surface of the vasculature. Th1 chemokines are essential for the recruitment of CXCR3^+^ cytolytic T-cells and NK cells, whereas VCAM1 and ICAM1 mediate T-cell adhesion and diapedesis. However, not all blood vessels are functionally equivalent in their expression/display of CAMs or chemokines. Instead, these factors predominate in HEVs, post-capillary venules (PCVs)/veins, and in the less well-characterized “IFN ECs” that are enriched in the expression of IFN signature genes, including *Cxcl9/10*, and appear in sites of local inflammation (23). Thus, therapeutically controlling or shaping the plasticity of the tumor vasculature offers an intriguing solution to boost anti-tumor immunity and augment ICB efficacy. Understanding the genetic or epigenetic mechanisms that mediate this plasticity will inform new strategies for altering blood vessel specialization in ways that drive anti-tumor immune cells into the TIME (24).

Since the epigenetic mechanisms that control EC functions in tumors are largely unknown, in this study, we used a conditional deletion approach in ECs to investigate how targeting the epigenetic factor DNA methyltransferase 1 (Dnmt1) impacts angiogenesis, tumor progression, and modulation of anti-tumor immunity. Knocking out Dnmt1 in ECs using in vitro and in vivo (*Dnmt1*^iECKO^) models potentiates striking activation of IFNγ/TNFα-driven pathways, boosts recruitment of memory CD4^+^ T-cells that are required for tumor suppression, and augments ICB in a model of experimental lung metastases. These results were especially profound using melanoma cells with high tumor mutational burden, suggesting an intriguing link between tumor cell mutational heterogeneity and the epigenetic mechanisms that generate immune-permissive tumor vasculature.

## RESULTS

### *Dnmt1*^iECKO^ impairs tumor growth and angiogenesis in non-immunogenic tumors, whereas highly immunogenic tumors regress

To evaluate the impact of *Dnmt1*^iECKO^ (Cdh5^Cre^:Dnmt1^fl/fl^:ZSGreen^l/s/l^) on tumor growth and angiogenesis, we implanted two isogenic melanoma cell lines, YUMM1.7 (low immunogenicity) and YUMMER1.7 (high immunogenicity), into control or *Dnmt1*^iECKO^ mice (25). We noticed that distinct growth patterns emerged depending on tumor immunogenicity. YUMM1.7 tumors displayed rapid growth in control mice (Cdh5^Cre^:ZSGreen^l/s/l^) but showed a reduced rate of growth in *Dnmt1*^iECKO^ mice (Fig. 1, a-b). However, ∼ 60% of YUMMER1.7 tumors regressed in *Dnmt1*^iECKO^ mice, with occasional mice showing no tumor burden at the time of euthanasia (Fig. 1, c-d). We hypothesized that reduced angiogenesis in YUMMER1.7 tumors could account for these differences. To test this hypothesis, we analyzed tumor-associated angiogenesis using binary and skeletonized imaging to quantify vascular structures using whole tumor scans. Using three different parameters (number of vessel branches, branch length, and number of vessel junctions), these data revealed significant reductions in angiogenesis in both tumor types implanted in control and *Dnmt1*^iECKO^ mice (Fig. 1, e-g). However, no statistically significant differences in angiogenesis were found in either tumor type implanted in *Dnmt1*^iECKO^ mice, suggesting alternative mechanisms may account for the regression of YUMMER1.7 tumors; including, for example, alterations in the composition of the TIME.

**Figure 1.**
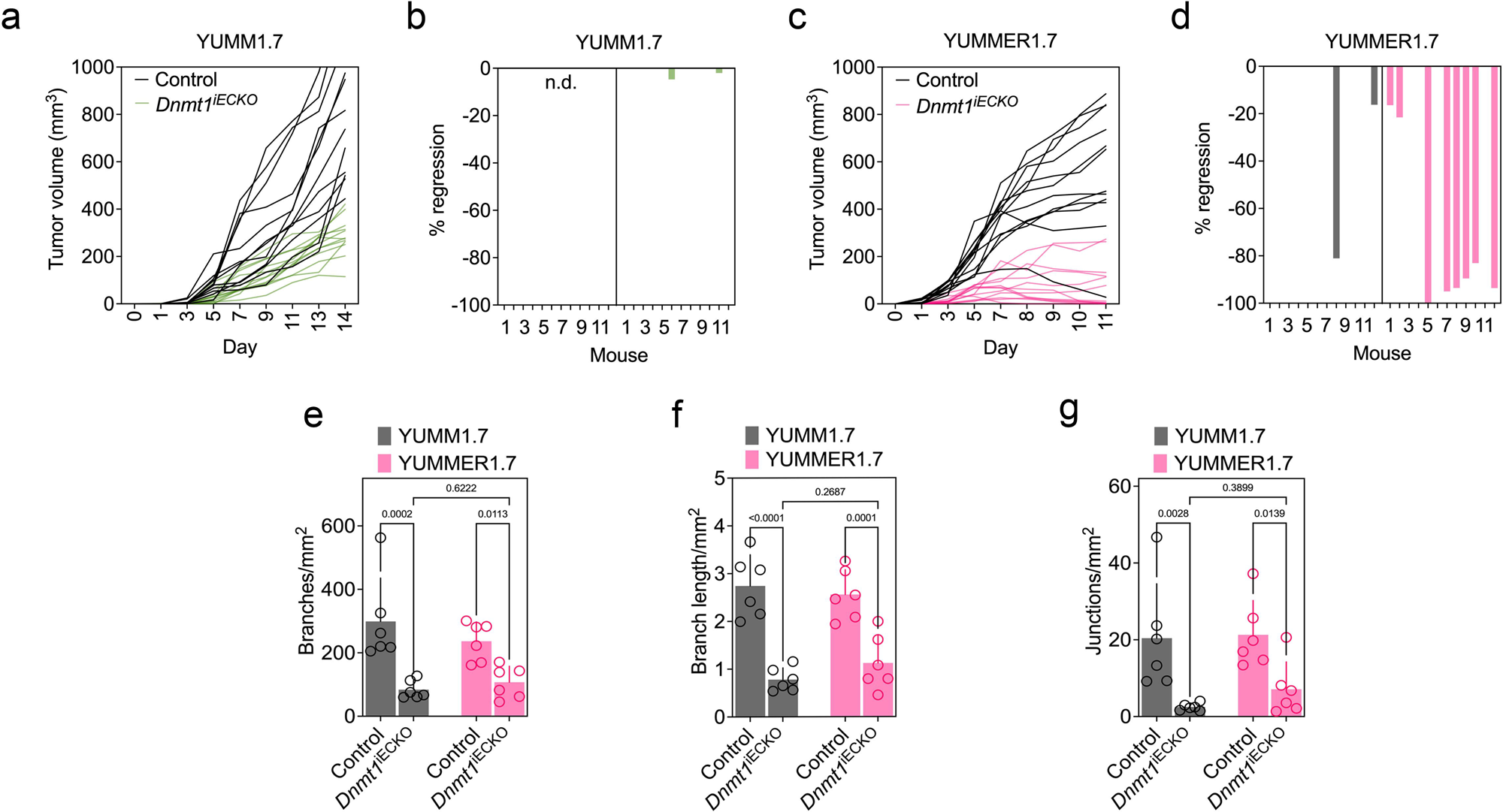
*Dnmt1*^iECKO^ impairs tumor growth and angiogenesis in non-immunogenic tumors, whereas highly immunogenic tumors regress. (a-d) Growth of YUMM1.7 and YUMMER1.7 tumors subcutaneously inoculated in control versus *Dnmt1*^iECKO^ mice. Tumor volumes were measured with calipers every other day. Data analysis was performed using a two-way ANOVA (*n*=12 mice per group). (e-g) The total numbers of vessel branches, branch lengths, and branch junctions per tumor was quantified using ImageJ. Statistical analysis was conducted using a two-way ANOVA (*n*=6 tumors per group).

### *Dnmt1*^iECKO^ reshapes the tumor immune microenvironment, which includes increases in memory CD4^+^ and CD8^+^ T cells, vessel-associated T-cells, and VCAM1^+^ vasculature

To investigate the impact of *Dnmt1*^iECKO^ on the TIME, we compared and contrasted selected immune cell populations using FACS in collagenase-dispersed YUMM1.7 and YUMMER1.7 tumors (26). No differences were observed in the fractions of myeloid-lineage cells or regulatory T-cells between the two tumor types (Supplemental Fig. 2, a-c). However, NK cells, while no different in control versus *Dnmt1*^iECKO^ mice in YUMM1.7 tumors, were increased by ∼2-fold in YUMMER1.7 tumors in *Dnmt1*^iECKO^ mice (Supplemental Fig. 2, d). To our surprise, depleting NK cells, while slightly increasing tumor growth in YUMMER1.7 tumors, did not rescue the growth of YUMMER1.7 tumors in *Dnmt1*^iECKO^ mice (Supplemental Fig. 2, e-f). These findings suggest that different immune cells, such as CD4⁺ or CD8⁺ T cells, may be responsible for tumor regression in YUMMER1.7 tumors in *Dnmt1*^iECKO^ mice.

In contrast to myeloid-lineage cells or regulatory T-cells, significant alterations in T-cell subset populations were observed between control and *Dnmt1*^iECKO^ tumors. Notable increases in the frequency of naive CD4^+^ T cells and central memory CD4^+^ and CD8^+^ T-cells were found in YUMMER1.7 tumors in *Dnmt1*^iECKO^ mice when compared to control tumors (Fig. 2, a-d). We next performed immunofluorescence staining in these tumors to determine the spatial organization of infiltrated T-cells. These data revealed increased CD3^+^ T-cell infiltration in both tumor types implanted in *Dnmt1*^iECKO^ mice, but a more pronounced 2-4 increase in total CD3^+^ T-cells and a doubling of vessel-associated T-cells observed in YUMMER1.7 tumors in *Dnmt1*^iECKO^ mice (Fig. 2, e-g). Since VCAM1 is a well-known adhesion molecule that binds to α4β1 on T-cells, we stained these tumors with anti-VCAM1 antibodies. These data showed significantly increased (2-4 fold) VCAM1^+^ vessel densities, especially in YUMMER1.7 tumors (Fig. 2, h-j). These findings demonstrate that *Dnmt1*^iECKO^ changes the composition of the TIME, including increased infiltration of memory T-cells and an activated tumor vasculature characterized by upregulated VCAM1, especially in tumors with a high tumor mutational burden. Collectively, these changes could account for the regression of YUMMER1.7 tumors in *Dnmt1*^iECKO^ mice.

**Figure 2.**
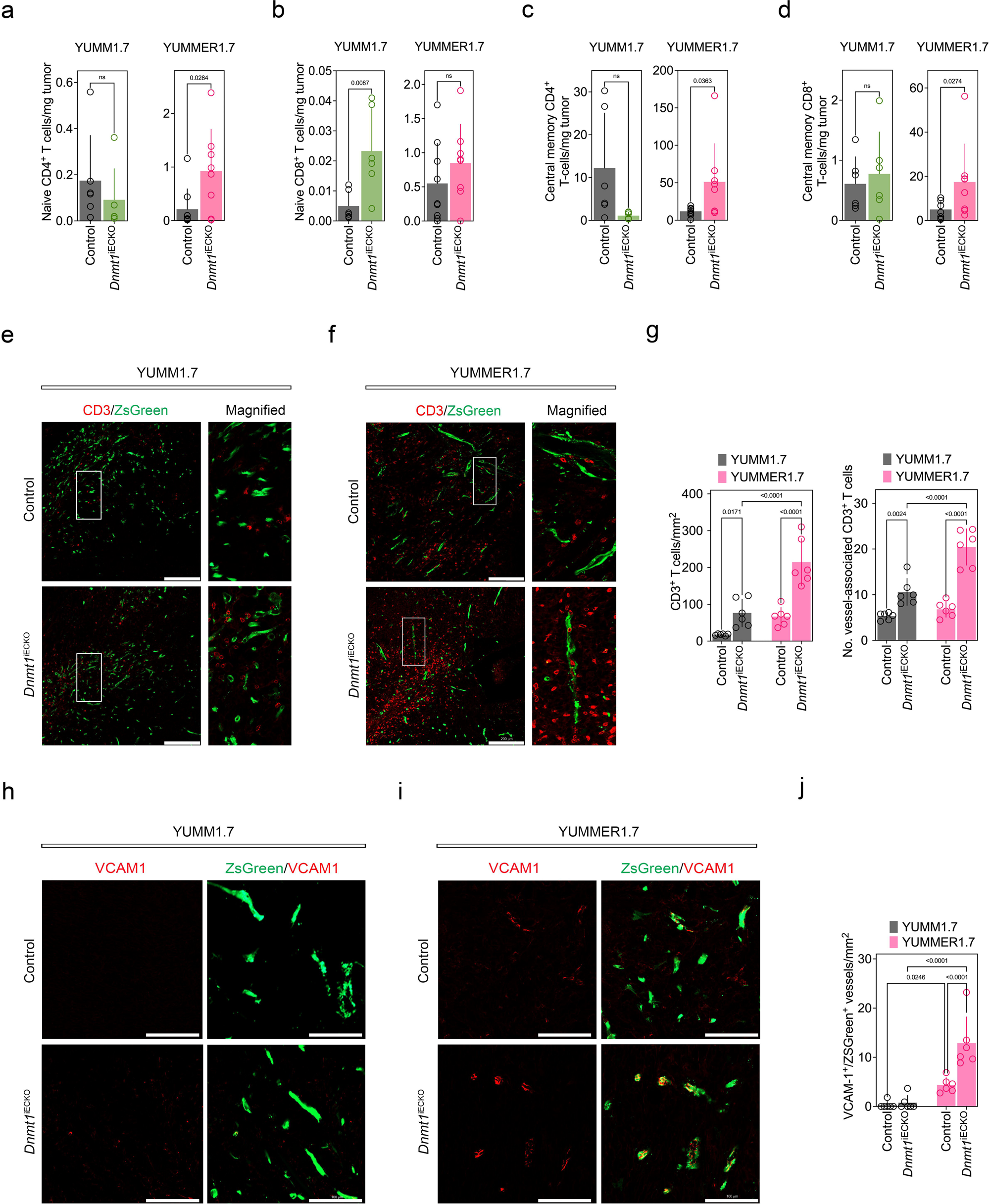
*Dnmt1*^iECKO^ reshapes the tumor immune microenvironment, which includes increases in memory CD4^+^ and CD8^+^ T cells, vessel-associated T-cells, and VCAM1^+^ vasculature. (a-d) Flow cytometric quantification of CD4 and CD8 T-cell subsets in YUMM1.7 versus YUMMER1.7 tumors from control versus *Dnmt1*^iECKO^ mice. Panels depict the numbers of naïve CD4^+^ T-cells, naïve CD8^+^ T-cells, central memory CD4^+^ T-cells, and central memory CD8^+^ T-cells. Statistical analyses were performed by the Mann-Whitney test. (e-f) Representative histology for control and *Dnmt1*^iECKO^ tumors stained with anti-CD3 antibody. Insets indicate CD3^+^ T-cells near ZSGreen^+^ blood vessels. Scale bar, 200 μm. (g) Quantitative analysis of total CD3^+^ T-cells and CD3^+^ T-cells associated with blood vessels. (h-i) Immunostaining for VCAM1 (red), alongside ZSGreen-labeled blood vessels (green). Scale bar, 100 μm. (j) Quantification confirms increased VCAM1-expressing vasculature in YUMMER1.7 tumors in *Dnmt1*^iECKO^ mice compared to controls. For all quantitative analyses in this series, each data point is an individual mouse.

### Tumors in *Dnmt1*^iECKO^ mice show increased granzyme B^+^ cells and cleaved caspase-3 alongside augmented anti-tumor immunity

To investigate the mechanism whereby highly immunogenic tumors regress in *Dnmt1*^iECKO^ mice, we carried out immunostaining for granzyme B (Grz B) which is a marker for cytotoxic T-cells. These data showed a significantly enhanced infiltration of Grz B^+^ immune cells within *Dnmt1*^iECKO^ tumors, particularly in YUMMER1.7 tumors, relative to respective controls (Fig. 3a and Supplemental Fig. 3). Notably, Grz B^+^ lymphocytes were larger and appeared elongated in YUMMER1.7 tumors, suggesting they might be more motile or activated in the context of *Dnmt1* deletion in the vasculature (Fig. 3,b-c). The pattern of Gz B positivity associated with higher numbers of cleaved caspase 3^+^ (CC3) cells in the TIME (Fig. 3d and Supplemental Fig. 3). These findings indicate that *Dnmt1*^iECKO^ enhances the recruitment and activation of cytotoxic lymphocytes and promotes robust apoptosis in highly immunogenic melanoma.

**Figure 3.**
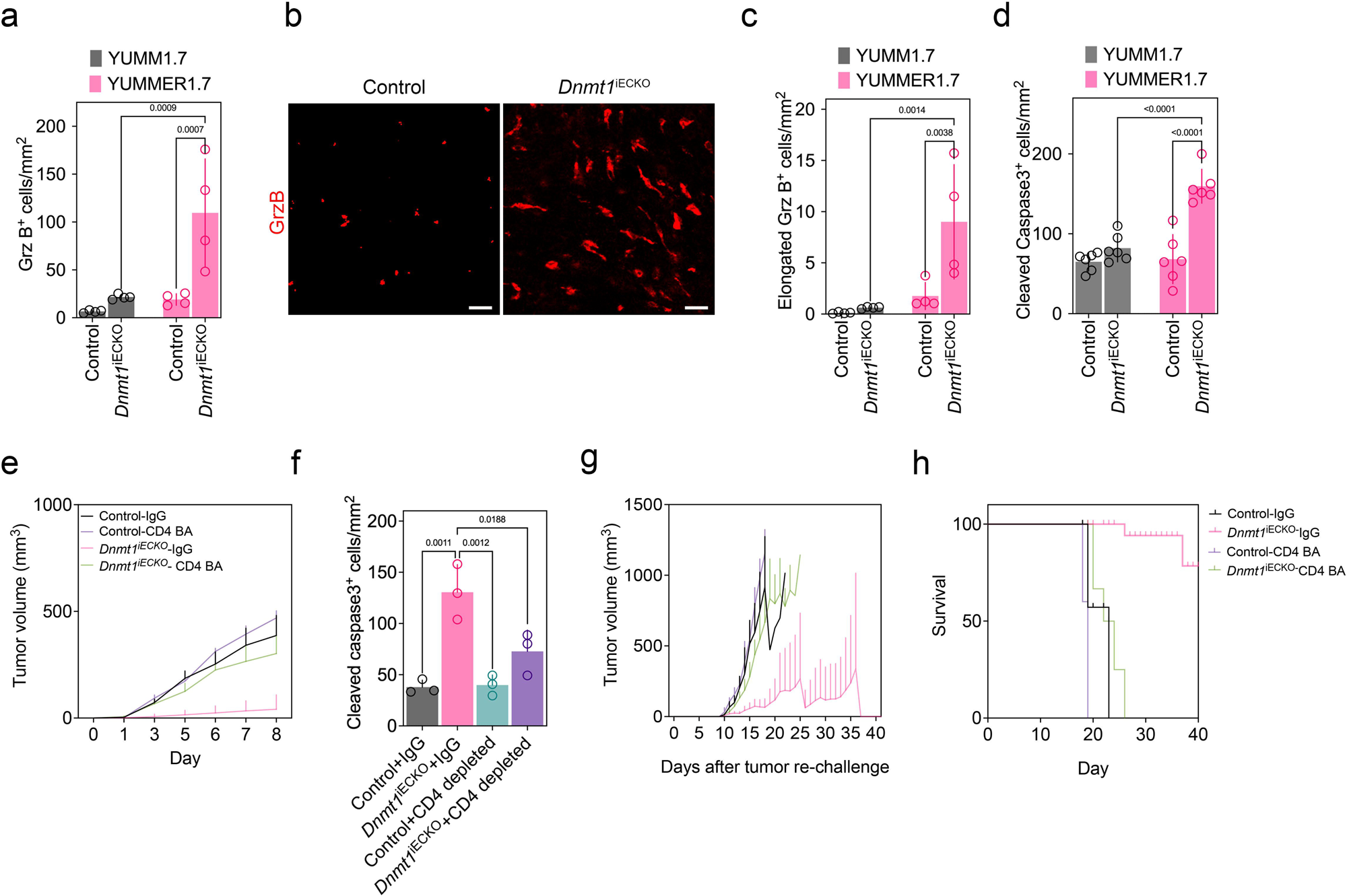
Tumors in *Dnmt1*^iECKO^ mice show increased granzyme B^+^ cells and cleaved caspase-3 alongside augmented anti-tumor immunity. (a) Quantification of total Grz B^+^ lymphocytes in control versus *Dnmt1*^iECKO^ mice. (b) Representative immunohistochemical staining of Granzyme B (Grz B; red) and DAPI (grey) in YUMMER1.7 tumors in control versus *Dnmt1*^iECKO^ mice showing the elongated phenotype. (c) Quantification reveals significant increases in elongated Grz B^+^ cells in YUMMER1.7 tumors in *Dnmt1*^iECKO^ mice compared to controls. Scale bar, 100 μm. Data from n=4 mice were analyzed using a two-way ANOVA. (d) Quantification of CC3^+^ cells in tumors inoculated in control versus *Dnmt1*^iECKO^ mice. (e) Growth curves for YUMMER1.7 tumors in control versus *Dnmt1*^iECKO^ mice following CD4 T-cell depletion *(n*=5 mice per group). (f) Quantification of CC3^+^ cells in CD4^+^ T-cell-depleted mice. (g) Growth curves for YUMMER1.7 tumors in control versus *Dnmt1*^iECKO^ mice after tumor re-challenge and CD4 T-cell depletion. The legend in “g” is the same as in “e”. (h) Kaplan-Meier survival plots for YUMMER1.7 tumors in control versus *Dnmt1*^iECKO^ mice after tumor re-challenge and CD4 T-cell depletion. For all quantitative analyses in this series, each data point is an individual mouse.

Our previous work showed an important role for CD8^+^ T-cells for mammary tumor suppression in *Dnmt1*^iECKO^ mice. In the present study, we found an unexpected but robust increase in naive CD4^+^ and central memory CD4^+^ T-cells in YUMMER1.7 tumors in *Dnmt1*^iECKO^ mice. To investigate the role of CD4^+^ T-cells in mediating the anti-tumor immune response observed in *Dnmt1*^iECKO^ melanoma models, we used a CD4 T-cell blocking antibody (CD4 BA) to deplete CD4^+^ T cells in control or *Dnmt1*^iECKO^ mice. In contrast to NK cell depletion, depleting CD4^+^ T-cells enhanced tumor growth and survival in *Dnmt1*^iECKO^ mice, as did blocking the egress of lymphocytes from the lymph node using FTY-720 (Fig. 3e and Supplemental Fig. 3). To analyze effector function after CD4 T-cell depletion, we again examined tumors for CC3 positivity. Depleting CD4^+^ T-cells reduced CC3^+^ cells in *Dnmt1*^iECKO^ mice in contrast to IgG controls (Fig. 3f). Since CD4^+^ memory T-cells were elevated in *Dnmt1*^iECKO^ mice, we carried out additional depletions of CD4^+^ T-cells followed by tumor resection/rechallenge studies in control versus *Dnmt1*^iECKO^ mice. To establish T-cell memory, control and *Dnmt1*^iECKO^ mice were initially challenged with 1 × 10^5^ YUMMER1.7 cells. Tumors were then surgically resected three days after injection. After tumor removal, mice underwent a secondary challenge with 1 × 10^7^ cells, following CD4^+^ T-cell depletion. In control mice, tumor growth after rechallenge was comparable between IgG-treated control and CD4^+^ T-cell BA-treated *Dnmt1*^iECKO^ mice (Fig. 3g). However, tumor growth was enhanced following CD4^+^ T-cell depletion in *Dnmt*1^iECKO^ mice, whereas 3/5 IgG-treated *Dnmt1*^iECKO^ mice demonstrated tumor rejection and 2/5 showed tumor growth after rechallenge (Fig. 3g). Consequently, *Dnmt1*^iECKO^ mice exhibited prolonged survival following tumor rechallenge compared to controls (Fig. 3h). Taken together, these data suggest that CD4^+^ T cells are critical for driving an anti-tumor immune response of immunogenic tumors in *Dnmt1*^iECKO^ mice, but not all mice necessarily form functional memory T-cells evidenced by a lack of tumor rejection upon tumor rechallenge.

### Immune checkpoint blockade is more effective against experimental lung metastases in *Dnmt1*^iECKO^ mice, as deleting Dnmt1 in ECs provokes robust potentiation of cytokine stimulation and up-regulation of vascular co-stimulatory molecules

Cancer immunotherapies are typically used in patients with metastatic disease. Thus, we tested combinations of anti-PDL1 and anti-CTLA4 in a model of experimental metastases to lung using control versus *Dnmt1*^iECKO^ mice. After validating the efficiency of tamoxifen-induced, EC-specific deletion of *Dnmt1*, we treated mice with luciferase-tagged YUMMER1.7 cells via the tail vein (Fig. 4a). After 10 days, mice were treated i.p. with the drug combinations. After 21 days, bioluminescence imaging was performed. The results show that Dnmt1 deletion in the vasculature was sufficient to suppress tumor burden in the lung, with 5/10 mice showing minimal to no tumor burden (Fig 4. b-c). Combinations of anti-PDL1 and anti-CTLA4 were also potent at suppressing lung metastases, with 7/10 mice showing tumor burden but at a significantly reduced size compared to controls. In *Dnmt1*^iECKO^ mice treated with the drug combination, only 2/10 mice showed evidence of tumor burden whereas 8/10 mice were tumor-free. In *Dnmt1*^iECKO^ mice that formed tumors, we found that combinations of ICB resulted in significantly greater numbers of CD3^+^ T-cells compared to *Dnmt1*^iECKO^ alone (Fig. 4d). These data suggest that combinations of ICB, in the context of *Dnmt1* deletion in endothelium, result in increased numbers of CD3^+^ T-cells in the lung TME and improved tumor control.

**Figure 4.**
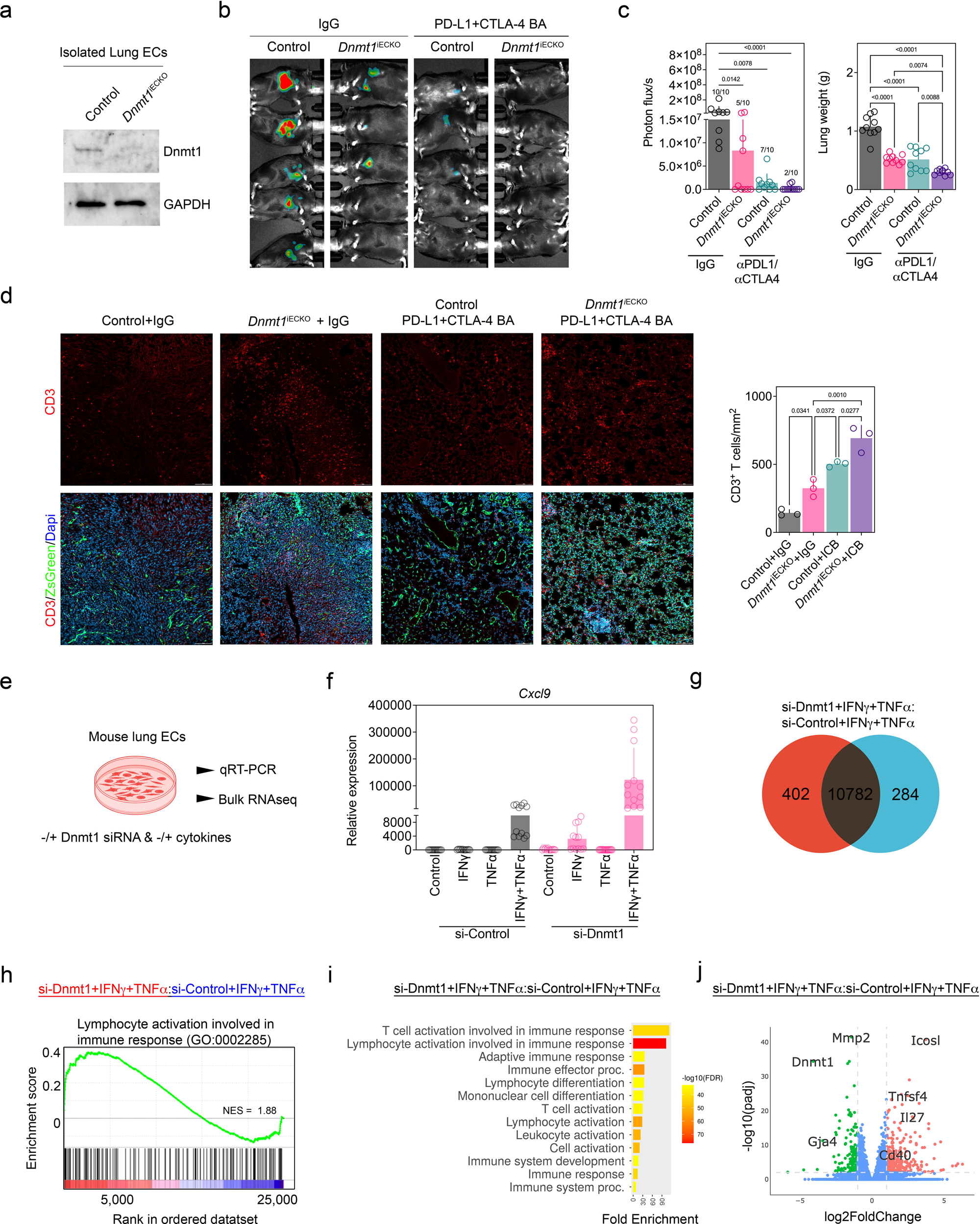
Immune checkpoint blockade is more effective against experimental lung metastases in *Dnmt1*^iECKO^ mice, as deleting *Dnmt1* in ECs provokes robust potentiation of cytokine stimulation and up-regulation of vascular co-stimulatory molecules. (a) Western blot for Dnmt1 protein levels in CD31^+^/CD45^-^/ZsGreen^+^ ECs sorted by FACS. (b) Representative bioluminescence images of luciferase-tagged YUMMER1.7 tumors in the indicated mice. (c) Photon influx was measured by Lago X imaging and lung weights were determined at the end of the study. Data were analyzed using a Kruskal-Wallis test (photon flux) or ANOVA (lung weights). Treatments included IgG, anti-PD-L1, anti-CTLA4, or a combination of anti-PD-L1/CTLA4 blockade in control versus *Dnmt1*^iECKO^ mice. (d) Immunofluorescence analysis of metastatic lung tissues, stained for CD3 antibody (red) and quantified. Blood vessels are ZSGreen^+^. Scale bar, 100 μm. For all quantitative analyses in this series, each data point is an individual mouse. (e) Schematic for bulk RNAseq in mouse lung ECs treated with Dnmt1 siRNA −/+ cytokines. RNAseq samples were run in quadruplicate. (f) qRT-PCR analysis for the candidate gene *Cxcl9* in these EC cultures under the indicated treatment. Each data point is an individual sample. (g) Venn diagram representing unique genes in the indicated samples. (h) GSEA showing enrichment of genes for lymphocyte activation in the immune response (GO:0002285) in the indicated samples. (i) Core enrichment genes from the GSEA were further analyzed using ShinyGO 0.8, and the significantly-enriched pathways were plotted. (j) Volcano plot depicting significantly up-(orange) and down-regulated genes (green) in the indicated samples. Several genes important for T-cell co-stimulation were enriched in ECs treated with Dnmt1-siRNA + IFNγ/TNFα and are indicated on the plot.

Our previous work showed that deleting Dnmt1 in ECs primes the vasculature for stimulation by IFNγ or TNFα, resulting in upregulation of selected CAMs and Th1 chemokines (26). These data are consistent with activation of type I IFN responses following Dnmt1 inhibition in other contexts (27, 28). To determine in an unbiased way how deletion of Dnmt1 in ECs impacts responses to these cytokines, we silenced Dnmt1 in mouse lung ECs, followed by stimulation with IFNγ, TNFα, or the combination (Fig. 4e). Before bulk RNAseq, we validated the expression of the known IFN-inducible gene, Cxcl9, by qRT-PCR. These data showed the expected potentiation in Cxcl9 expression following si-Dnmt1 and IFNγ stimulation and a robust augmentation in Cxcl9 expression following treatment with both cytokines simultaneously, even in control samples (Fig. 4f). These data are consistent with a strong synergy between these cytokines in the stimulation of IFNγ and TNFα response patterns in ECs.

Since both IFNγ and TNFα are present in the microenvironment of most tumors, we next sought to determine how silencing Dnmt1 in ECs might globally alter gene expression when both cytokines are operative. Using bulk RNAseq in these EC cultures, we found a total of 402 up-regulated and 284 down-regulated genes when *Dnmt1* was silenced alongside IFNγ/TNFα in comparison to IFNγ/TNFα-treated ECs (Fig. 4g). Notably, Gene Set Enrichment Analysis (GSEA) showed a trending, although not statistically significant, signature corresponding to lymphocyte activation involved in immune response (Fig. 4h). Further exploration of the genes enriched in this pathway using GO Enrichment analysis showed multiple pathways important for anti-tumor immunity including T-cell activation, adaptive immune responses, and lymphocyte differentiation (Fig. 4i). Interestingly, silencing *Dnmt1* alongside IFNγ/TNFα resulted in up-regulation of several co-stimulatory molecules in ECs; in particular, those that are important for driving T-cell memory such as *Icosl*, *Tnfsf4*, and *Cd40* (Fig. 4j). These data correspond with the increased numbers of memory T-cell subsets we observed in *Dnmt1*^iECKO^ mice in vivo and suggest that epigenetically altering the tumor vasculature is sufficient to change the composition of the TIME.

### Single-cell RNAseq of tumor-associated ECs in *Dnmt1*^iECKO^ mice reveals a phenotypic shift in the vasculature, including greater numbers of post-capillary venule ECs, venous ECs, and IFN-ECs

To assess how endothelial-specific *Dnmt1* loss influences TEC heterogeneity, we performed scRNA sequencing of CD31⁺/CD45⁻ TECs from control versus *Dnmt1*^iECKO^ tumors and compared the dataset with a reference lung EC atlas containing normal lung ECs (NECs) and TECs (29). UMAP visualization after integration and clustering identified the expected EC subtypes, including capillary-1, capillary-2, proliferating ECs, arterial and venous ECs, post-capillary venule (PCV) ECs, lymphatic ECs, interferon-stimulated ECs (IFN-ECs), activated-artery ECs, and TEC-capillary populations (Fig. 5a). These populations were consistently present in both control and *Dnmt1*^iECKO^ samples, showing that EC *Dnmt1* loss does not disrupt global EC lineage identity. Comparison of NECs and TECs within the reference dataset revealed distinct TEC-specific phenotypes that were largely absent in NECs (Fig. 5b). NEC-derived clusters are shown in yellow tones, while TEC-derived clusters are shown in blue tones. As expected, *Dnmt1* expression was markedly decreased in *Dnmt1*^iECKO^ TECs (Fig. 5c). By contrast, key EC markers such as *Cdh5* (VE-Cadherin) and *Pecam1* remained consistent across both control and *Dnmt1*^iECKO^ TECs, indicating preserved endothelial marker expression. Quantification of cluster abundance showed modest shifts in EC subtypes between the samples. Notably, tumors from *Dnmt1*^iECKO^ mice showed increased proportions of venous ECs (∼2-fold, p adj=1.25e^-6^), PCV ECs (∼2-fold, p adj=1.28e^-4^), and IFN-stimulated ECs (∼1.5-fold, p adj =7.44e^-2^) (Fig. 5d). To our surprise, GSEA analysis did not show a statistically significant enrichment for global IFNγ-, IFNα-, and TNFα-related pathways in TECs from *Dnmt1*^iECKO^ mice; however there were trending increases in these pathways in TECs from *Dnmt1*^iECKO^ mice (Supplemental Fig. 4). This may be due to the small total numbers of TECs in the IFN-ECs cluster. While IFN-ECs are a less well-characterized population of ECs in general, PCVs are major sites of lymphocyte infiltration across the vasculature, due to their increased expression of CAMs such as *Vcam1*. Indeed, examination of a candidate T-cell attracting chemokine (*Cxcl9*) and *Vcam1*, showed they were enriched in PCVs and IFN-ECs in *Dnmt1*^iECKO^ mice (Fig. 5e). Interestingly, *Vcam1* appeared elevated in additional subpopulations of TECs in *Dnmt1*^iECKO^ mice, including tip cells, proliferating ECs, and activated artery ECs. Since IFN response pathways were trending upward in TECs from *Dnmt1*^iECKO^ mice, we examined two additional core enrichment genes, including *Cd40* and *Cd274,* and found they were modestly elevated in IFN-ECs and PCV-ECs. Thus, the tumor vasculature in *Dnmt1*^iECKO^ shows a modest shift in vascular specification with a trend towards greater numbers of PCV and IFN ECs. Since these types of ECs are specialized for lymphocyte entry, this could account for the improved efficacy of ICB in this setting.

**Figure 5.**
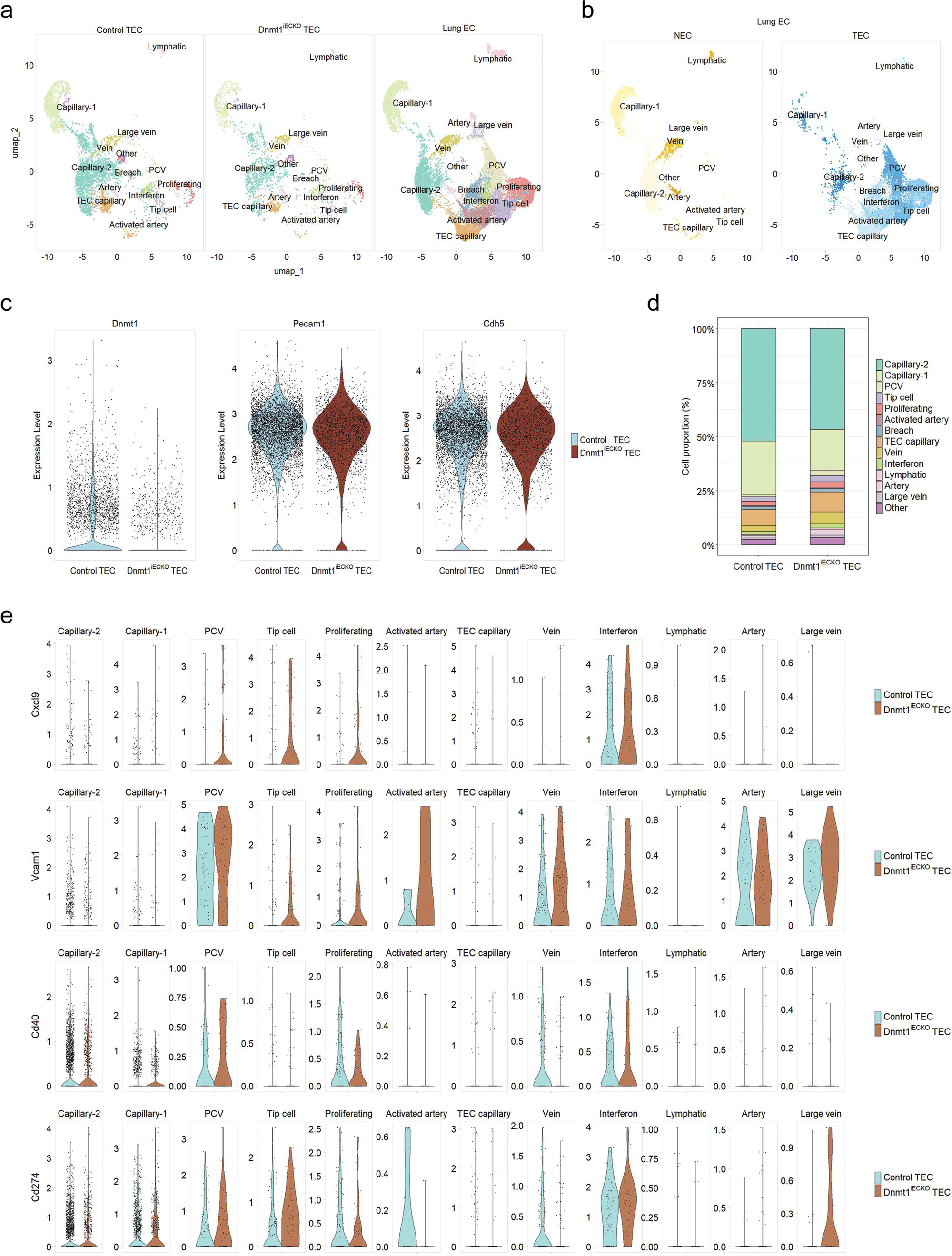
Single cell RNAseq of tumor-associated ECs in *Dnmt1*^iECKO^ mice reveals a phenotypic shift in the vasculature, including greater numbers of post-capillary venule ECs, venous ECs, and IFN-ECs. **(a)** UMAP visualization of integrated CD31⁺/CD45⁻ tumor endothelial cells (TECs) from control and *Dnmt1*^iECKO^ tumors, showing major endothelial subtypes. (b) Reference lung EC atlas depicting normal ECs (NECs; yellow tones) and TECs (blue tones). (c) Violin plots showing reduced *Dnmt1* expression in *Dnmt1*^iECKO^ TECs, whereas EC markers *Cdh5* and *Pecam1* are consistently expressed across samples. (d) Proportion plot of TEC subtypes in control and *Dnmt1*^iECKO^ tumors. (e) Cluster-level differential gene expression analysis.

## DISCUSSION

Anti-tumor immune cells, such as cytotoxic T-cells or NK cells, must have access to the tumor microenvironment for active immune surveillance and for cancer immunotherapies to be effective. To do so, these types of immune cells must first adhere to ECs, roll along the surface, and then extravasate across the vasculature. This process is coordinated, in part, by CAMs and T-cell-attracting chemokines that are displayed on the surface of the endothelium. In the tumor vasculature, these factors are often aberrantly expressed or downregulated, which impairs immune cell mobility and extravasation (30–33). Thus, reprogramming the vasculature with judicious doses of anti-angiogenic drugs or other therapies to reinforce CAM/chemokine expression has the potential to augment the efficacy of cancer immunotherapies (34–37).

Our previous findings demonstrated that in mice with conditional deletion of *Dnmt1* in the vasculature, the expression of CAMs and T-cell-attracting chemokines was elevated, which potentiated ICB efficacy in mammary tumor models (26). In the current study, we have further defined a role for vascular *Dnmt1* in coordinating anti-tumor immunity using both lowly immunogenic and highly immunogenic melanoma cells. Our data, in brief, show that *Dnmt1*^iECKO^ mediates direct and indirect anti-tumor effects, including the induction of immunomodulatory ECs, potentiation of IFNγ- and TNFα-driven genetic programs, promotion of CD4⁺ T-cell memory responses, and augmentation of adaptive immunity. Consequently, combining *Dnmt1*^iECKO^ with ICB therapy significantly suppressed lung metastasis in mice, which was associated with greater numbers of CD3^+^ T-cells in the lung. These outcomes may be due to changes in the composition and phenotype of the tumor vasculature, including increased proportions of PCVs and IFN-ECs in *Dnmt1*^iECKO^ mice. IFN-ECs are a poorly understood subpopulation of ECs that assemble in sites of inflammation and in tumors, and they appear to be important for anti-tumor immunity (38, 39). It is possible that IFN-ECs are assembled from activated PCV and that, due to the potentiating effect on IFN signaling that is driven by *Dnmt1* inhibition, the transition between PCVs and IFN-ECs is facilitated in *Dnmt1*^iECKO^ mice.

Some limitations of our study include the realization that, in the EC conditional deletion model used herein, *Dnmt1* is disabled in all ECs and thus not restricted to the tumor vasculature. Therefore, systemic deletion of *Dnmt1* in other organ or tissue microenvironments (e.g. lymph nodes) could account for the anti-tumor immune responses we observed. To circumvent this, targeted deletion of Dnmt1 using vascular-tropic AAVs, or other means, would be needed (40). Similarly, the scRNAseq results did not necessarily phenocopy the bulk RNAseq data showing expression of multiple co-stimulatory molecules in pure EC cultures stimulated with cytokines. But the different kinetics of short-term in vitro stimulation versus long-term in vivo stimulation where there are confounding heterotypic cell-cell interactions occurring, could account for this discrepancy. Furthermore, while flow cytometry data and antibody depletion studies indicate an important role for CD4^+^ memory T-cells during tumor growth inhibition in *Dnmt1*^iECKO^ mice, we cannot say with certainty whether the CD4^+^ T-cells are the source for cytolytic granzymeB production in this study (41). Our previous work in mammary tumor models showed an important role for CD8^+^ T-cells during the inhibition of mammary tumor growth in *Dnmt1*^iECKO^ mice. But the cell lines used in this study were not of the low-versus high TMB dichotomy. It is therefore possible that vascular Dnmt1, while shaping anti-tumor immunity, does so via different immune-mediated mechanisms in different contexts. Finally, based on our bulk RNAseq data from in vitro cultures and scRNAseq in vivo, both immune-stimulating (e.g. *Icosl* and *Cd40*) and immune-inhibitory (e.g. *Cd274* and *Lgals9*) factors are elevated in ECs when *Dnmt1* is targeted alongside pro-inflammatory cytokines. This likely represents one of the many usual feedback mechanisms that orchestrate the immune response to pathogens or tumors (42). Nevertheless, based on our study, the net effect of *Dnmt1* inhibition in ECs, at least in the short-term, is augmented anti-tumor immunity. It is possible that over time, the immune-inhibitory signals elicited by targeting *Dnmt1* in the vasculature could eventually lead to tolerance and eventual tumor escape from immune destruction.

Anti-cancer immunity encompasses the surveillance, detection, and elimination of neoplastic cells, which relies on the essential process of immune cells infiltrating the EC barrier to mount an effective immune response (43). Growing evidence suggests that immunomodulatory ECs play a pivotal role in enhancing T-cell activation and boosting anti-tumor immunity, often in reciprocity with multiple immune cell subsets (18, 23, 44–46). Our study reveals that modulating an epigenetic switch (i.e. Dnmt1) in ECs is sufficient to change tumor vascular phenotype in ways that convert the TIME from immune-restrictive to immune-permissive.

## MATERIALS AND METHODS

### Reagents and cell lines

Antibodies used in this study were anti-mouse DNMT1 (Abcam), anti-mouse GAPDH, anti-mouse CD8 (Biolegend, BioXcell, or eBiosciences), anti-mouse CD4 (BD Biosciences and BioXcell), anti-mouse NK1.1 (BioXcell), anti-mouse IgG (BioXcell), and anti-mouse PD-L1 (Genentech, MTA program). MDEC (mouse skin EC) and Veravec (mouse lung ECs) were cultured in 1 g/l D-glucose DMEM (LG-DMEM) with 10% FBS, 10% Nu Serum IV, 10 ng/ml VEGF, 5 ng/ml bFGF, and 100 mg/l porcine heparin (47–50). YUMM1.7, YUMMER1.7, or YUMMER1.7-luciferase cells were cultured in 4.5 g/l D-glucose DMEM and 10% FBS. Cells were maintained at 37°C in a 5% CO_2_ atmosphere supplemented with 20% O_2_.

### Animal models

The following mouse strains were used: C57BL/6 wild-type mice and transgenic mice on the C57BL/6 background, *VEcad*-Cre^ERT2^ (Rha);*ZsGreen^l/s/l^* mice (used as control mice) were crossed with *Dnmt1*^fl/fl^ mice to generate *VECad*-Cre^ERT2^;*Dnmt*1^fl/fl^;*ZsGreen^l/s/l^*mice.

### Quantitative real-time polymerase chain reaction

RNA was extracted from tissues or ECs using the RNA purification kit (Zymo Research). cDNA synthesis was carried out using an iScript synthesis kit (Bio-Rad). Quantitative Reverse Transcription PCR (qRT-PCR) was performed with SYBR Green (Invitrogen). Data were normalized to the GAPDH control. The results were analyzed via the ΔΔCt method.

### Immunohistochemistry

Harvested tumor tissue was placed in 4% PFA in PBS for 24 hours at 4°C. Tumors were then transferred to 30% sucrose in PBS for cryoprotection. After sucrose, tumors were embedded in OCT. Serial tissue sections were performed at −20 °C using a sliding microtome. Sections were fixed with acetone and were washed in 1× PBS/0.2% Triton X-100 before blocking with 10% normal goat serum, 4% bovine serum albumin in 1× PBS/0.2% Triton X-100 for 1 hr. Primary antibodies were applied overnight for cryosections. After washing in 1× PBS/Triton X-100, the secondary antibody was used for 2 hrs. After nuclei staining with DAPI (100 ng/mL in 1×PBS; Molecular Probes) for 30 min, sections were embedded in Vectashield (Molecular Probes).

### Image processing and morphological analysis

Vessel length analysis was performed using Fiji by applying a threshold transformation that maximizes the global average contrast of vessel edges. After binary images were obtained, they were processed as follows: the blood vessel structure was first extracted from the background. Then, skeletonization was applied to the binary images. The lengths of the blood vessel branches and the branch numbers were then measured.

### Flow cytometry

For endothelial cell sorting, tumor tissues from control and *Dnmt1*^iECKO^ mice were chopped into small pieces and incubated for 90 min at 37 °C in dissociation buffer containing 2 mg/ml of collagenase type I (Worthington), 1 mg /ml of DNase (Worthington), and 2.5 units/ml of neural protease (Worthington). The digested tissues were separated into a single-cell suspension by passing through a 100 μm cell strainer. The cells were stained with primary antibodies for 30 minutes on ice, followed by incubation with secondary antibodies. Samples were fixed with 2% paraformaldehyde. Cells were then analyzed using flow cytometry analysis on a FACS Caliber apparatus (BD) and FlowJo software (Tree Star Inc.) to quantify cell populations.

### In vivo tumor models

For the primary tumor growth experiment, mice were injected subcutaneously into the shaved shoulder with YUMM1.7 or YUMMER1.7 melanoma cells suspended in 100 μl of PBS, containing 5 x 10^5^ cells. Mice were monitored for the appearance of tumors after injection to begin caliper measurements. For the tumor rechallenge experiment, an initial 1 x 10^5^ cells of YUMMER1.7 were injected into control and *Dnmt1*^iECKO^ mice. Palpable tumors were surgically removed and rechallenged with 1 x 10^7^ cells of YUMMER1.7. The mice were treated with CD4 blocking antibody (400 mg/kg) or isotype-matched IgG control antibody. For the depletion experiments, IgG anti-NK1.1 (10 mg/kg, Clone PK136, BioXCell), anti-CD4 (20 mg/kg, Clone GK1.5, BioXCell), or anti-CD8a (20 mg/kg, Clone 2.43, BioXCell) antibodies were administered by i.p. injection. Anti-NK1.1 antibody was injected into the mice every 2 days for the duration of the experiment. For the depletion of CD4 T-cells, antibodies were administered 7, 10, 13, and 15 days after tumor inoculation. In the experimental lung metastasis study, 5 × 10^5^ YUMMER1.7-luciferase-expressing cells in 100 μl of PBS were injected via the tail vein. After 1 week, the mice were treated with anti-PD-L1 (10 mg/kg, Genentech MTA program) and CTLA-4 (5 mg/kg, 9D9, BioXCell) antibodies every four days. On day 21, the mice were monitored for bioluminescence and euthanized, and the lung tumors were examined further.

### Generation of YUMMER1.7 expressing luciferase stable cell line and bioluminescence imaging

The pLV-lenti construct was co-transformed into 293T cells with pMD2.G and psPAX2 constructs using Lipofectamine 2000 transfection reagent. Lentiviral supernatants were harvested at 72 hrs post-transfection and filtered through a 0.45-μm membrane. YUMMER1.7 cells were infected for 48 hrs with fresh lentivirus with 8 μg/ml polybrene and cultured for 48 hrs. The activity of luciferase was analyzed using a luminometer (PerkinElmer). To image YUMMER1.7^Luc^ in vivo, mice were intraperitoneally injected with 100 μl of Cycluc1 (Sigma Aldrich). The animals were anesthetized with isoflurane (2% in 1 l/min O_2_), and bioluminescence images were acquired using the LagoX (Spectral Instrumental Imaging). Images were acquired every 2 mins for 30 mins total (10 s exposure/image). Images were analyzed using Aurora software. Regions of interest (ROIs) were drawn around each cell mass, and the total number of photons within each ROI was recorded.

### Preparation of ECs for bulk RNA sequencing

Mouse lung ECs were seeded at 1×10^5^ cells/mL in 6-well plates and cultured overnight at 37 °C in a humidified 5% CO_2_ incubator. The next day, cells were transfected with either control siRNA or DNMT1-targeting siRNA (BLOCK-iT oligos; Invitrogen) using Lipofectamine RNAiMAX (Thermo Fisher Scientific) in Opti-MEM I Reduced Serum Medium for 4 h, following the manufacturer’s instructions. A BLOCK-iT Alexa Fluor Red fluorescent oligo (Invitrogen) was used in parallel wells to monitor transfection efficiency. After 4 hrs, the transfection medium was replaced with growth medium containing 20% serum, and the cells were cultured for 24 hrs at 37 °C, humidified 5% CO₂. A second siRNA transfection was performed 24 h later to reinforce the knockdown. Cells were then treated with TNFα (10 ng/mL) or IFNγ (1,000 U/mL) for 16 hrs. Total RNA was isolated using a column-based RNA extraction kit (Zymo Research) according to the manufacturer’s instructions. Libraries were prepared and sequenced by Novogene for bulk RNA-seq analysis. Experiments were performed with *n* = 4 biological replicates per condition.

### FTY720 treatment

Vehicle control (DMSO) and FTY720 (Cayman Chemical) were administered via oral gavage at 1 mg/kg. Treatment with FTY720 began seven days after the inoculation of tumor cells, and it was administered daily throughout the entire study duration. Tumor dimensions were recorded each day using calipers, with the measurements performed in a blinded manner. At the end of the experiment, the animals were euthanized, and tumor tissues were harvested for analysis.

### Tumor endothelial cell (TEC) isolation and sorting

Tumors (∼1 cm³) from three control mice and three Dnmt1^iECKO^ mice were harvested, and tumors within each sample were pooled to obtain sufficient EC numbers. Samples were collected in ice-cold low-glucose DMEM (LG-DMEM), washed thoroughly, and minced into fragments <5 mm under sterile conditions. Tissue fragments were transferred to GentleMACS C tubes containing a digestion mix of collagenase II (2 mg/mL, Worthington LS004176), dispase (2.5 U/mL, Worthington LS02104), and DNase I (1 mg/mL, Worthington LS002006). Tumors were digested at 37 °C for 30–45 min with agitation (120 rpm) and processed on a GentleMACS Dissociator using program *m_imptumor_02* to achieve single-cell suspensions.

Digested samples were passed through 100µm strainers, washed with FACS buffer (PBS + 0.5% BSA + 2 mM EDTA), and centrifuged at 1200 rpm for 5 min. Red blood cell lysis was performed using 1× Pharm Lyse B. Cell suspensions were first incubated with Live-or-Dye viability dye (Biotium 32002), followed by Fc block (Miltenyi 130-092-575) for 10 min on ice. Cells were then stained with PE-conjugated CD31 (BD 553373) and APC-conjugated CD45 (BD 559864) for 30 min on ice.

Live CD31⁺CD45⁻ tumor endothelial cells (TECs) were sorted on an Influx cell sorter (Becton Dickinson) into LG-DMEM supplemented with 10% FBS, washed, and resuspended in PBS + 0.04% BSA. Final cell suspensions were adjusted to 500–700 cells/µL with >85% viability. Sorted TECs were immediately used for 10x Genomics Chromium Single Cell 3’ v3 library preparation according to the manufacturer’s protocol.

### Single-cell RNA-seq library preparation and sequencing

Sorted cells were processed using the 10x Genomics Chromium Single Cell 3′ Gene Expression kit (GEM-X v4) following the manufacturer’s protocol. Libraries were prepared targeting ∼10,000 cells per sample and sequenced on an Illumina NextSeq 2000 using the P2 XLEAP-SBS 100-cycle kit (paired-end) at a depth of 20,000–25,000 reads per cell.

### 10x cloud data processing

Raw FASTQ files were processed using 10x Genomics Cloud Analysis with Cell Ranger Multi v9.0.1. Libraries were specified as GEM-X 3′ Gene Expression v4 and aligned to the mouse reference genome (mm10, 2020-A). The pipeline generated filtered gene–barcode matrices for downstream single-cell analysis.

### Computational analysis of scRNA-seq data

Raw gene–barcode matrices from control and Dnmt1^iECKO^ TEC samples were processed using Seurat v5. Low-quality cells with high mitochondrial content (>10%) or low gene complexity (<9000 detected genes) were removed. Control and *Dnmt1*^iECKO^ TECs were evaluated for quality control, and doublets were identified using DoubletFinder v3. Only singlet cells were retained for downstream analysis. To obtain a high-purity TEC dataset, cells expressing immune (Ptprc), epithelial (Epcam), pericyte (Cspg4), or smooth muscle (Myh11) markers were excluded. Endothelial identity was confirmed based on expression of Cdh5 and/or Pecam1. After all filtering steps, the final dataset included 4,977 control TECs and 2,075 Dnmt1^iECKO^ TECs.

A publicly available lung endothelial reference dataset provided as normalized count and metadata CSV files was imported into Seurat to create the reference object. This dataset originally contained NEC, TEC, and additional treatment-associated EC populations. Only the NEC and TEC subsets were retained and aligned to their metadata and included in the integration workflow solely to support EC subtype annotation. It was not used for differential comparisons between control and *Dnmt1*^iECKO^ TECs.

To integrate the datasets, the control TECs, *Dnmt1*^iECKO^ TECs, and lung EC reference subset were combined using Seurat’s integration workflow to allow joint visualization and cluster identification based on shared gene features. Clustering was performed at a resolution of 0.5. UMAP embeddings were generated for the visualization of major EC populations. Cluster-defining marker genes were identified using Seurat’s differential expression analysis, and the top markers were used to annotate endothelial subclusters for downstream interpretation.

### Statistics

All values are expressed as ± standard error of the mean (SEM). Results were analyzed using a student’s t-test or ANOVA using GraphPad Prism 10 software. For tumor volumes and weights, the significance level was determined using ANOVA, Sidek’s multiple comparisons tests (volumes), or a 2-tailed Student’s t-test (weights). P values less than 0.05 were considered significant.

## Data availability

Source data and code will be provided upon publication

## Acknowledgements

ACD is supported by grants from the the National Institutes of Health/National Cancer Institute (2RO1 CA177875 and RO1 CA2558451). Portions of this research were supported by the NCI Cancer Center Support Grant 5P30CA044579 and by the UVA Genome Analysis and Technology Core (RRID:SCR_018883). Additional support was provided by The University of Virginia Flow Cytometry Core (RRID: SCR_017829). The Sony MA900 Cell Sorter was funded through the NIH S10 instrument program (S10 Grant Number 1S10OD028518-1).

## Author contributions statement

DJK and ACD conceptualized the study and wrote the manuscript. DJK and YS carried out experiments and data analysis. MM and MR carried out flow cytometry analysis of tumors. SA and CR analyzed scRNAseq and bulk RNAseq data, respectively. All authors have been provided with a copy of the complete manuscript prior to submission.

## Supplementary information

**Supplementary figure 1. Blood vessel densities are reduced in both YUMM1.7 and YUMMER1.7 tumors in *Dnmt1*^iECKO^ mice.** (a,b) Representative binary and skeletonized images highlight differences in the density and organization of tumor-associated vasculature. A confocal tile scan captured entire tumors in control and *Dnmt1*^iECKO^ mice. Tumor blood vessels were visualized using ZsGreen labeling and converted into an 8-bit binary image (grey). The magnified boxed areas on the right allow for detailed observation and were inverted to emphasize lateral branches. Notably, arrowheads in control mice indicate branched vessels, contrasting with the straighter, narrower vessels in *Dnmt1*^iECKO^ mice. Scale bars = 1 mm.

**Supplementary figure 2. NK cells are increased in YUMMER1.7 tumors in *Dnmt1*^iECKO^ mice, but depleting NK cells does not rescue tumor growth.** (a-d) Flow cytometric quantification of myeloid-derived suppressor cells, regulatory T-cells, and NK cells in control versus *Dnmt1*^iECKO^ mice. Data analyzed using Student’s *t*-test. YUMMER1.7 growth curves following NK cell depletion. (e) Tumor-bearing mice were treated with anti-NK1.1 or isotype IgG starting from day zero. Tumor volumes were measured daily *(n* = 5 per group). No statistically significant differences were observed between the groups. (f) Tumor weights at endpoint from the same experiment shown in “e”, comparing control and *Dnmt1*^iECKO^ mice treated with IgG or anti-NK1.1 antibodies. For all quantitative analyses in this series, each data point is an individual mouse.

**Supplementary figure 3. GzB^+^ and CC3^+^ cells are increased in YUMMER1.7 tumors in *Dnmt1*^iECKO^ mice.** (a,b) Representative immunofluorescence images of tumor sections stained for GzB or CC3 (both shown in red) and counterstained with DAPI (grey). Scale bar = 200 μm. (c,d) Growth and overall survival of control versus *Dnmt1*^iECKO^ mice bearing YUMMER1.7 tumors and treated with FTY720.

**Supplementary figure 4. Enrichment scores in control versus *Dnmt1*^iECKO^ TECs.** (a) The plots showing trending ES for Hallmark TNFα, IFNγ, and IFNα response genes.

## References

1. A. C. Dudley, Tumor endothelial cells. Cold Spring Harb Perspect Med 2, a006536 (2012).

2. A. C. Dudley, A. W. Griffioen, Pathological angiogenesis: mechanisms and therapeutic strategies. Angiogenesis 26, 313–347 (2023).

3. E. Lanitis, M. Irving, G. Coukos, Tumour-associated vasculature in T cell homing and immunity: opportunities for cancer therapy. Nat Rev Immunol 10.1038/s41577-025-01187-w (2025).

4. D. M. Corey, Y. Rinkevich, I. L. Weissman, Dynamic Patterns of Clonal Evolution in Tumor Vasculature Underlie Alterations in Lymphocyte-Endothelial Recognition to Foster Tumor Immune Escape. Cancer Res 76, 1348–1353 (2016).

5. A. Bagaev et al., Conserved pan-cancer microenvironment subtypes predict response to immunotherapy. Cancer Cell 39, 845–865 e847 (2021).

6. J. Li et al., Remodeling of the immune and stromal cell compartment by PD-1 blockade in mismatch repair-deficient colorectal cancer. Cancer Cell 41, 1152–1169 e1157 (2023).

7. F. S. Hodi et al., TMB and Inflammatory Gene Expression Associated with Clinical Outcomes following Immunotherapy in Advanced Melanoma. Cancer Immunol Res 9, 1202–1213 (2021).

8. T. M. R. Noviello et al., Guadecitabine plus ipilimumab in unresectable melanoma: five-year follow-up and integrated multi-omic analysis in the phase 1b NIBIT-M4 trial. Nature Communications 14, 5914–5914 (2023).

9. P. A. Jones, H. Ohtani, A. Chakravarthy, D. D. De Carvalho, Epigenetic therapy in immune-oncology. Nat Rev Cancer 19, 151–161 (2019).

10. H. Y. Kuo, K. A. Khan, R. S. Kerbel, Antiangiogenic-immune-checkpoint inhibitor combinations: lessons from phase III clinical trials. Nat Rev Clin Oncol 21, 468–482 (2024).

11. D. Fukumura, J. Kloepper, Z. Amoozgar, D. G. Duda, R. K. Jain, Enhancing cancer immunotherapy using antiangiogenics: opportunities and challenges. Nat Rev Clin Oncol 15, 325–340 (2018).

12. M. Schmittnaegel et al., Dual angiopoietin-2 and VEGFA inhibition elicits antitumor immunity that is enhanced by PD-1 checkpoint blockade. Sci Transl Med 9 (2017).

13. S. Ragusa et al., Antiangiogenic immunotherapy suppresses desmoplastic and chemoresistant intestinal tumors in mice. J Clin Invest 130, 1199–1216 (2020).

14. E. J. M. Huijbers et al., Embryonic reprogramming of the tumor vasculature reveals targets for cancer therapy. Proc Natl Acad Sci U S A 122, e2424730122 (2025).

15. M. Schmittnaegel, M. De Palma, Reprogramming Tumor Blood Vessels for Enhancing Immunotherapy. Trends Cancer 3, 809–812 (2017).

16. F. Yang, G. Lee, Y. Fan, Navigating tumor angiogenesis: therapeutic perspectives and myeloid cell regulation mechanism. Angiogenesis 27, 333–349 (2024).

17. J. D. Peske, A. B. Woods, V. H. Engelhard, Control of CD8 T-Cell Infiltration into Tumors by Vasculature and Microenvironment. Adv Cancer Res 128, 263–307 (2015).

18. E. Allen et al., Combined antiangiogenic and anti-PD-L1 therapy stimulates tumor immunity through HEV formation. Sci Transl Med 9 (2017).

19. Y. Hua et al., Cancer immunotherapies transition endothelial cells into HEVs that generate TCF1(+) T lymphocyte niches through a feed-forward loop. Cancer Cell 41, 226 (2023).

20. M. Ramachandran et al., Tailoring vascular phenotype through AAV therapy promotes anti-tumor immunity in glioma. Cancer Cell 41, 1134–1151 e1110 (2023).

21. J. Verhoeven et al., Tumor endothelial cell autophagy is a key vascular-immune checkpoint in melanoma. EMBO Mol Med 15, e18028 (2023).

22. W. Ma et al., Targeting PAK4 to reprogram the vascular microenvironment and improve CAR-T immunotherapy for glioblastoma. Nat Cancer 2, 83–97 (2021).

23. J. Amersfoort, G. Eelen, P. Carmeliet, Immunomodulation by endothelial cells - partnering up with the immune system? Nat Rev Immunol 22, 576–588 (2022).

24. Y. Zhu, K. F. Brulois, T. T. Dinh, J. Pan, E. C. Butcher, COUP-TFII-mediated reprogramming of the vascular endothelium counteracts tumor immune evasion. Nat Commun 16, 7457 (2025).

25. J. Wang et al., UV-induced somatic mutations elicit a functional T cell response in the YUMMER1.7 mouse melanoma model. Pigment Cell Melanoma Res 30, 428–435 (2017).

26. D. J. Kim et al., Priming a vascular-selective cytokine response permits CD8(+) T-cell entry into tumors. Nat Commun 14, 2122 (2023).

27. M. L. Stone et al., Epigenetic therapy activates type I interferon signaling in murine ovarian cancer to reduce immunosuppression and tumor burden. Proc Natl Acad Sci U S A 114, E10981–E10990 (2017).

28. K. B. Chiappinelli et al., Inhibiting DNA Methylation Causes an Interferon Response in Cancer via dsRNA Including Endogenous Retroviruses. Cell 169, 361 (2017).

29. J. Goveia et al., An Integrated Gene Expression Landscape Profiling Approach to Identify Lung Tumor Endothelial Cell Heterogeneity and Angiogenic Candidates. Cancer Cell 37, 421 (2020).

30. D. M. Hellebrekers et al., Epigenetic regulation of tumor endothelial cell anergy: silencing of intercellular adhesion molecule-1 by histone modifications. Cancer Res 66, 10770–10777 (2006).

31. D. M. Hellebrekers et al., Identification of epigenetically silenced genes in tumor endothelial cells. Cancer Res 67, 4138–4148 (2007).

32. D. Vestweber, How leukocytes cross the vascular endothelium. Nat Rev Immunol 15, 692–704 (2015).

33. A. N. Woods et al., Differential Expression of Homing Receptor Ligands on Tumor-Associated Vasculature that Control CD8 Effector T-cell Entry. Cancer Immunol Res 5, 1062–1073 (2017).

34. Z. R. Huinen, E. J. M. Huijbers, J. R. van Beijnum, P. Nowak-Sliwinska, A. W. Griffioen, Anti-angiogenic agents - overcoming tumour endothelial cell anergy and improving immunotherapy outcomes. Nat Rev Clin Oncol 18, 527–540 (2021).

35. H. Yang et al., STING activation reprograms tumor vasculatures and synergizes with VEGFR2 blockade. J Clin Invest 129, 4350–4364 (2019).

36. A. H. Cleveland, Y. Fan, Reprogramming endothelial cells to empower cancer immunotherapy. Trends Mol Med 30, 126–135 (2024).

37. L. Wang-Bishop, et al., STING-activating nanoparticles normalize the vascular-immune interface to potentiate cancer immunotherapy. Sci Immunol 8, eadd1153 (2023).

38. L. S. Tombor et al., Immunoregulatory Endothelial Cells Interact With T Cells After Myocardial Infarction. Circ Res 137, 866–879 (2025).

39. S. L. Freshour et al., Endothelial cells are a key target of IFN-g during response to combined PD-1/CTLA-4 ICB treatment in a mouse model of bladder cancer. iScience 26, 107937 (2023).

40. M. Stamataki et al., Identification of AAV variants with improved transduction of human vascular endothelial cells by screening AAV capsid libraries in non-human primates. Gene Ther 10.1038/s41434-025-00563-4 (2025).

41. A. Cachot et al., Tumor-specific cytolytic CD4 T cells mediate immunity against human cancer. Sci Adv 7 (2021).

42. L. Chen, D. B. Flies, Molecular mechanisms of T cell co-stimulation and co-inhibition. Nat Rev Immunol 13, 227–242 (2013).

43. D. S. Chen, I. Mellman, Oncology meets immunology: the cancer-immunity cycle. Immunity 39, 1–10 (2013).

44. C. Qian, C. Liu, W. Liu, R. Zhou, L. Zhao, Targeting vascular normalization: a promising strategy to improve immune-vascular crosstalk in cancer immunotherapy. Front Immunol 14, 1291530 (2023).

45. R. Missiaen, M. Mazzone, G. Bergers, The reciprocal function and regulation of tumor vessels and immune cells offers new therapeutic opportunities in cancer. Semin Cancer Biol 52, 107–116 (2018).

46. B. Kruse et al., CD4(+) T cell-induced inflammatory cell death controls immune-evasive tumours. Nature 618, 1033–1040 (2023).

47. L. Xiao et al., Tumor Endothelial Cells with Distinct Patterns of TGFbeta-Driven Endothelial-to-Mesenchymal Transition. Cancer Res 75, 1244–1254 (2015).

48. L. Xiao, J. V. McCann, A. C. Dudley, Isolation and Culture Expansion of Tumor-specific Endothelial Cells. J Vis Exp 10.3791/53072, e53072 (2015).

49. J. V. McCann et al., Endothelial miR-30c suppresses tumor growth via inhibition of TGF-beta-induced Serpine1. J Clin Invest 129, 1654–1670 (2019).

50. A. C. Dudley et al., Calcification of multipotent prostate tumor endothelium. Cancer Cell 14, 201–211 (2008).

