## Supplementary figures and images for "An epigenetic switch in vascular phenotype augments anti-tumor immunity"

### s1

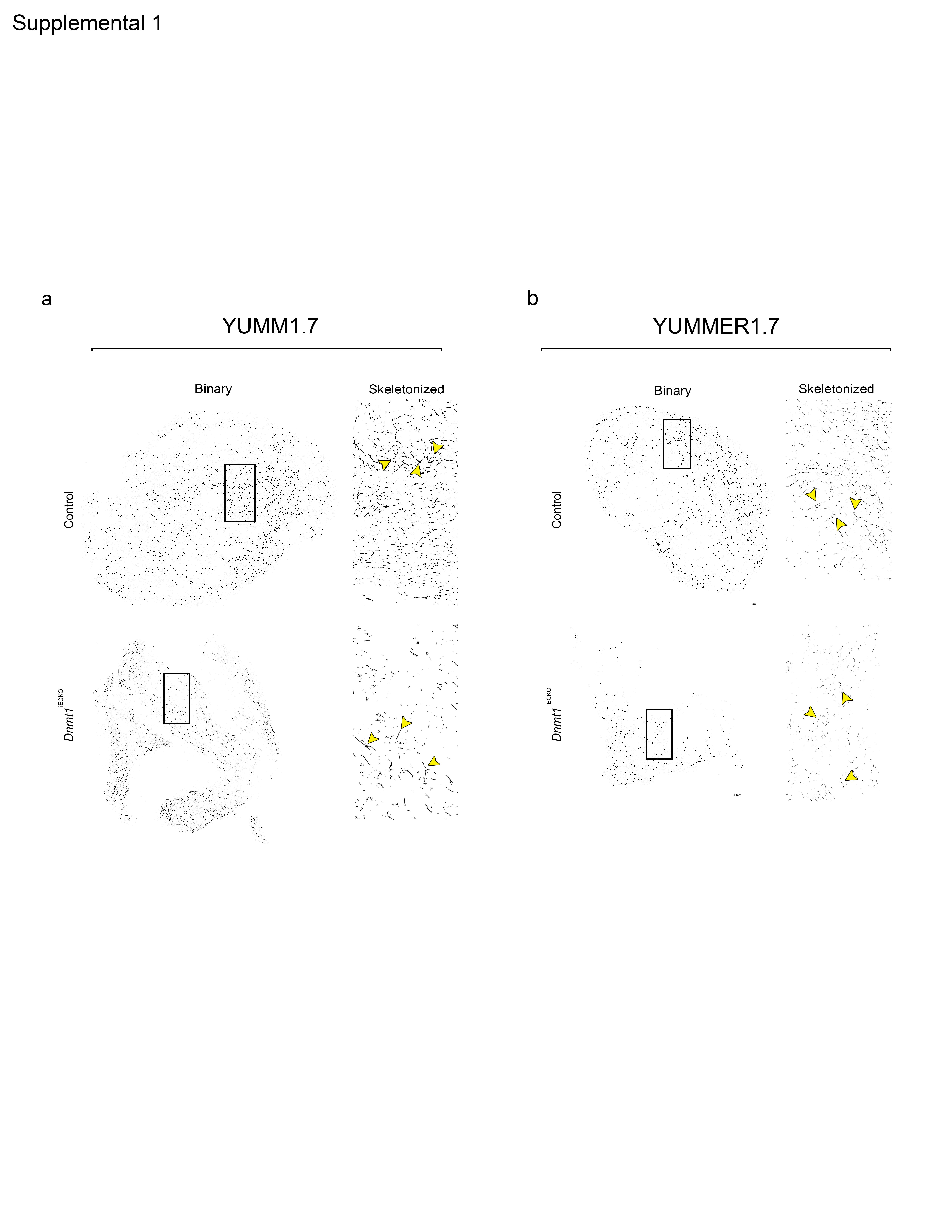

### s2

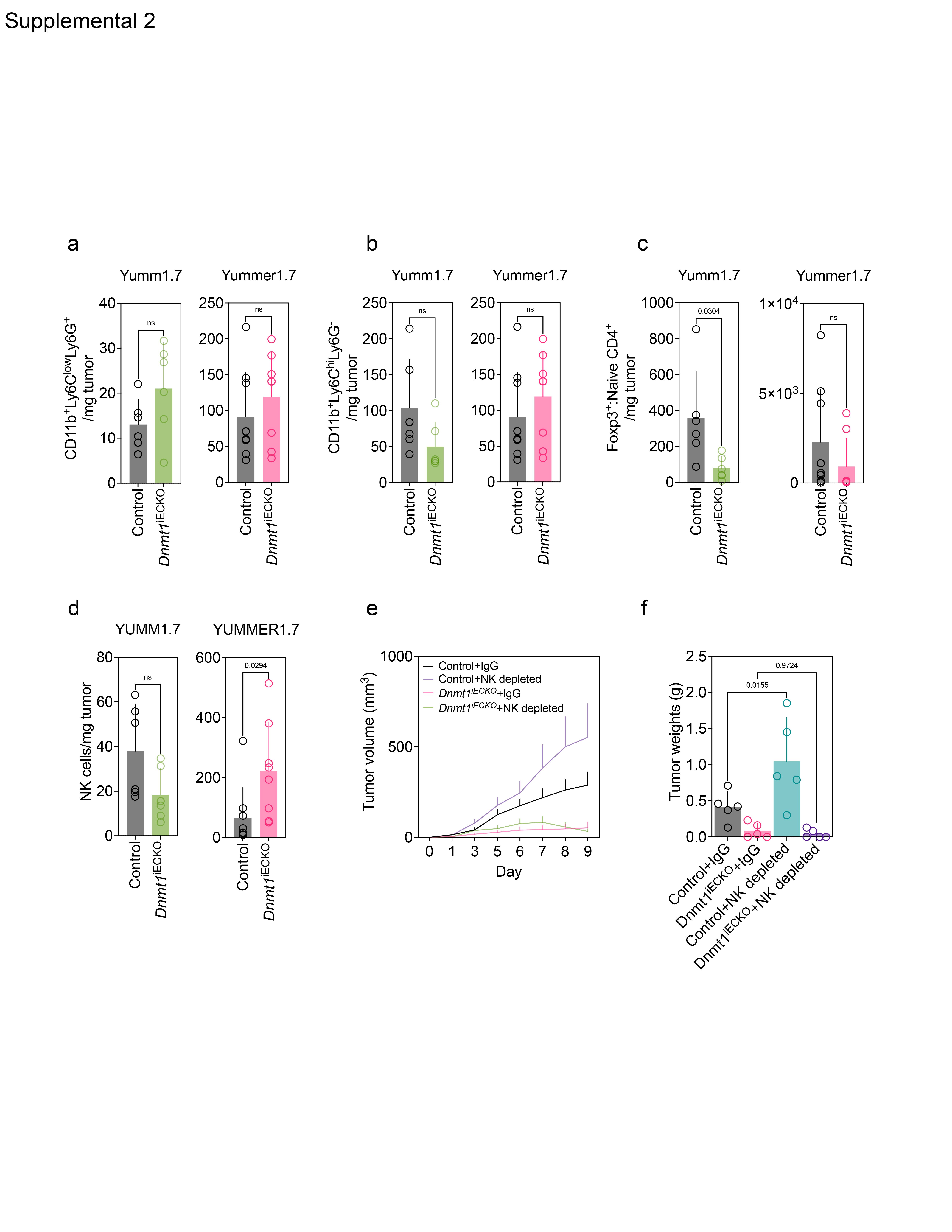

### s3

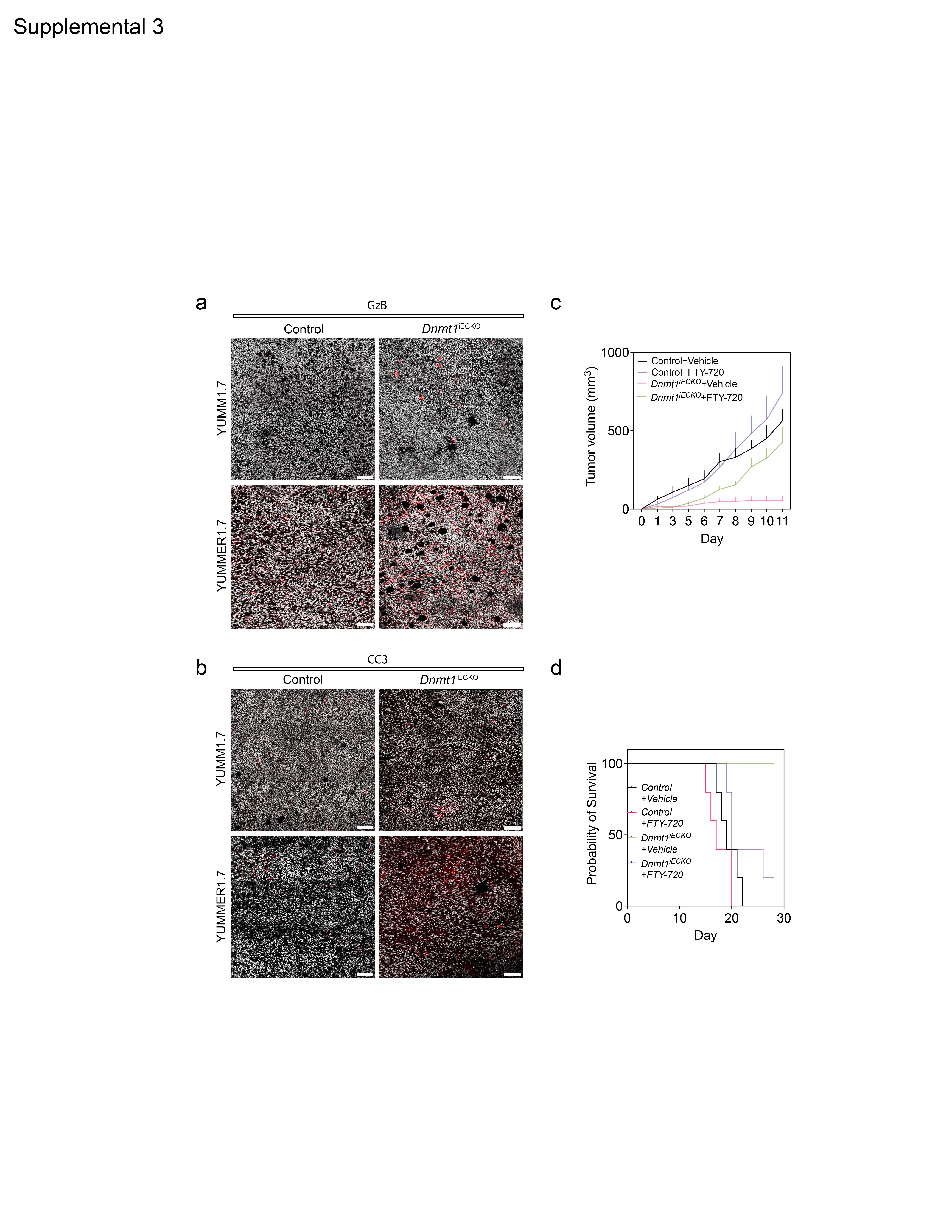

### s4

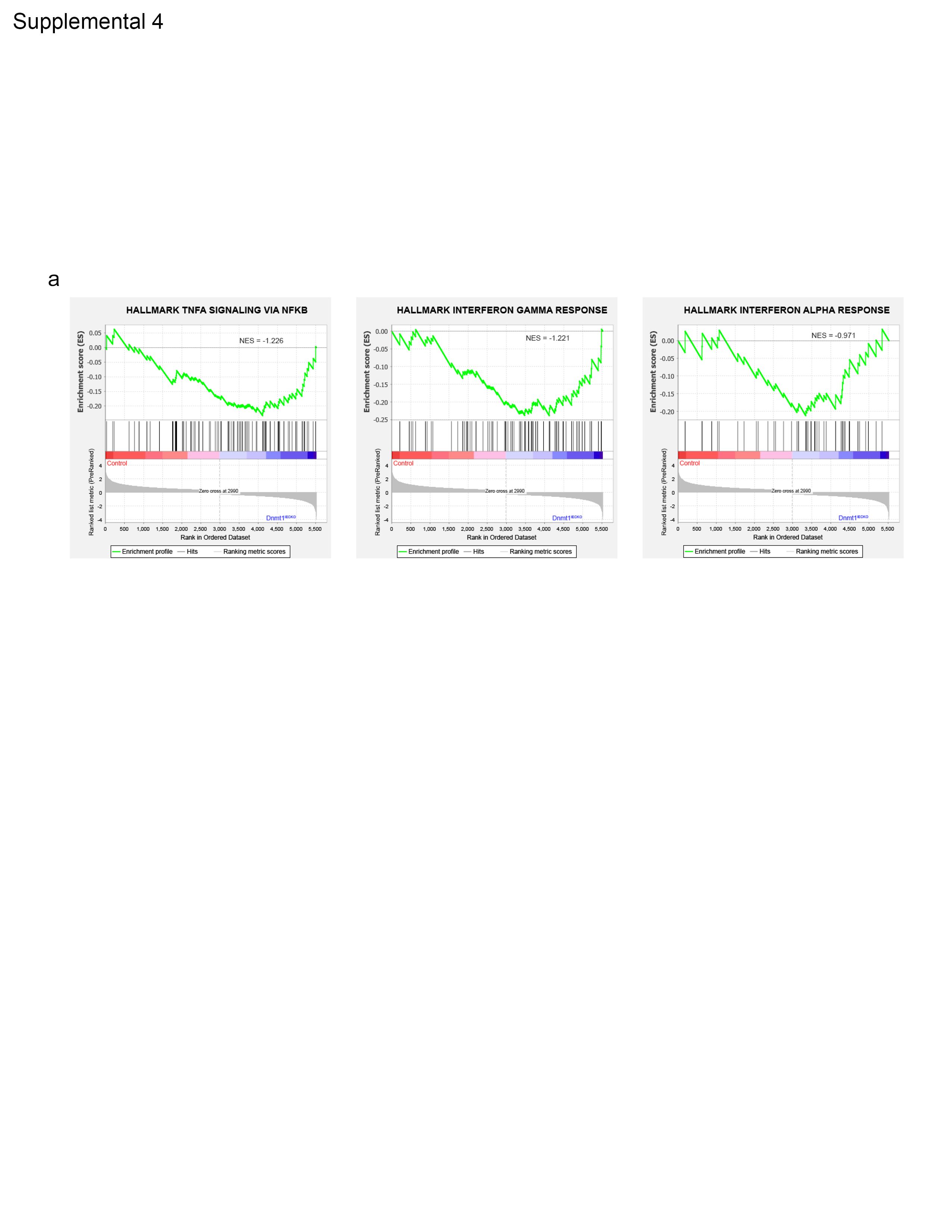
